# Select β-lactam combinations exhibit synergy against *Mycobacterium abscessus in vitro*

**DOI:** 10.1101/522649

**Authors:** Elizabeth Story-Roller, Emily C. Maggioncalda, Gyanu Lamichhane

## Abstract

*Mycobacterium abscessus* (*Mab*) is a nontuberculous mycobacterium that causes invasive pulmonary infections in patients with structural lung disease. *Mab* is intrinsically resistant to several classes of antibiotics and an increasing number of strains isolated from patients exhibit resistance to most antibiotics considered for treatment of *Mab* infections. Therefore, there is an unmet need for new regimens with improved efficacy to treat this disease. Synthesis of the essential cell wall peptidoglycan in *Mab* is achieved via two enzyme classes, L,D- and D-D-transpeptidases, with each class preferentially inhibited by different subclasses of β-lactam antibiotics. We hypothesized that a combination of two β-lactams that comprehensively inhibit the two enzyme classes will exhibit synergy in killing *Mab*. Paired combinations of antibiotics tested for *in vitro* synergy against *Mab* included dual β-lactams, a β-lactam and a β-lactamase inhibitor, and a β-lactam and a rifamycin. Of the initial 206 combinations screened, 24 pairs exhibited synergy. 13/24 pairs were combinations of two β-lactams. 12/24 pairs brought the minimum inhibitory concentrations of both drugs to within the therapeutic range. Additionally, synergistic drug pairs significantly reduced the frequency of selection of spontaneous resistant mutants. These novel combinations of currently-available antibiotics may offer viable immediate treatment options against highly-resistant *Mab* infections.

## INTRODUCTION

*Mycobacterium abscessus* (*Mab*) is considered to be among the most virulent of the rapidly-growing nontuberculous mycobacteria (NTM). It may be environmentally or nosocomially acquired (1) and can lead to severe and invasive pulmonary infections in immunocompromised patients or those with structural lung diseases, such as bronchiectasis or cystic fibrosis (CF). In the CF population, invasive *Mab* infections are associated with rapid lung function decline (2–4), so adequate treatment of these infections is paramount. *Mab* is intrinsically resistant to several classes of antibiotics and the percentage of clinical isolates exhibiting resistance to the few drugs currently available to treat this infection is steadily increasing (5–8). Sputum culture conversion rates as low as 25% have been described with antibiotic treatment alone (9) and the cure rate for *Mab* pulmonary disease is only 30-50% (10).

The current treatment guidelines for *Mab* pulmonary disease include at least 18 months of multi-drug therapy, several of which require intravenous administration and may be associated with significant cytotoxicity (11, 12). These recommendations are largely based on empirical evidence, as few systematic clinical trials have been performed to elucidate the optimal therapeutic regimen against *Mab*. It is also frequently necessary in clinical practice to tailor treatment regimens based on the resistance profiles of individual *Mab* isolates, as the high degree of variability in antibiotic resistance observed among *Mab* strains often precludes use of a standardized regimen (10).

Macrolide antibiotics have historically been considered the backbone of treatment against many NTMs, including *Mab* (2, 12). However, two of the three *Mab* subspecies, *abscessus* and *bolletii*, harbor a functional erm(41) gene, which confers inducible macrolide resistance and thus limits the effectiveness of this antibiotic class beyond the first two weeks of treatment (13, 14). Guidelines therefore recommend subspeciation of the *Mab* complex, which many clinical laboratories are not equipped to perform routinely. Consequently, some CF centers prescribe initial treatment regimens that include a combination of intravenous amikacin and either cefoxitin or imipenem rather than a macrolide (15). Cefoxitin (a cephalosporin) and imipenem (a carbapenem) are the only two β-lactam antibiotics included in the current *Mab* treatment guidelines (12, 16).

β-lactams function by inhibiting enzymes that catalyze synthesis of peptidoglycan, a three-dimensional macromolecule that forms the exoskeleton of bacterial cells (17). During the final step of peptidoglycan synthesis, *Mab* utilizes two enzyme classes; the canonical D,D-transpeptidases (DDT, also known as penicillin binding proteins) and the recently discovered L,D-transpeptidases (LDT) (18) to generate 4→3 and 3→3 linkages between stem peptides, respectively (Figure 1). Since as many as 80% of the linkages in *Mab* peptidoglycan are of the 3→3 type (18), the LDTs that generate them are likely at least as important as DDTs for this organism. An initial survey of the *Mab* genome identified five putative LDT encoding genes (19). These LDTs are differentially susceptible to β-lactam subclasses, with most carbapenems exhibiting strong inhibitory activities, followed by cephalosporins, and only a select few penicillins exhibiting moderate inhibition of these enzymes (20, 21).

**Figure 1.**
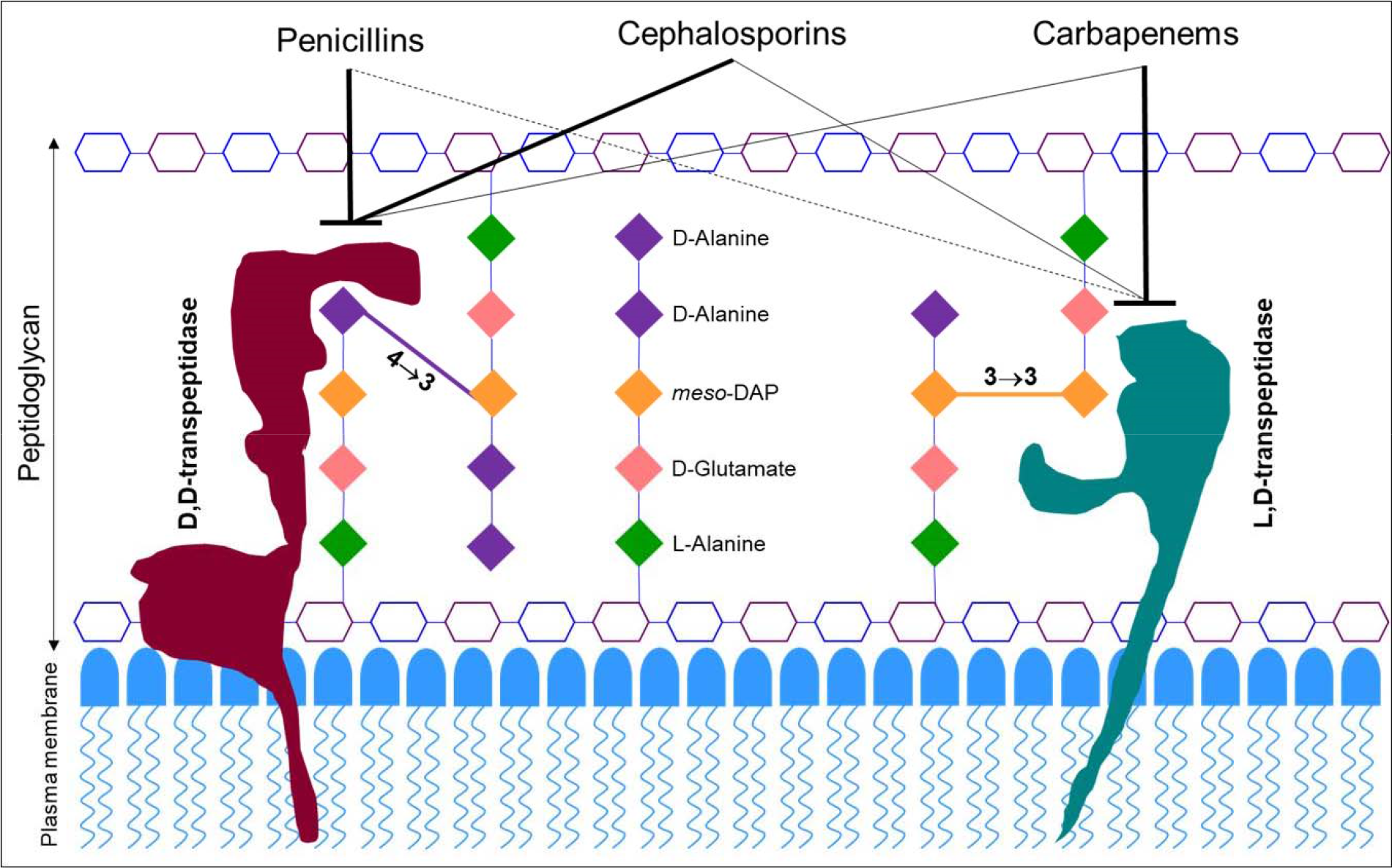
Model of *M. abscessus* peptidoglycan depicting preferential binding of β-lactam subclasses. The hexagonal structures represent sugars *N*-acetylglucosamine (purple) and *N*-acetylmuramic acid (blue).

Inhibition of peptidoglycan synthesis is lethal to bacteria (22). As LDT and DDT activities are required for synthesis of *Mab* peptidoglycan, simultaneous inhibition of both enzymes could be bactericidal. Since these enzymes exhibit differential susceptibilities to β-lactams (21, 23, 24), we hypothesize that a combination of β-lactam subclasses; one that optimally inhibits LDTs and another that specifically targets DDTs, will demonstrate synergy in killing *Mab*. In this study, we have tested this hypothesis by assessing potencies of combinatorial pairs against *Mab* using a panel of 16 β-lactams representing cephalosporins and carbapenems. Penicillins were not assessed, as many require frequent dosing and we preferentially chose oral cephalosporins requiring daily or twice daily dosing to simplify administration in patients.

*Mab* exhibits robust β-lactamase activity via Bla_Mab_, which significantly reduces the efficacy of β-lactams against this pathogen (25, 26). Bla_Mab_ degrades several β-lactams with much greater efficiency than BlaC of *M. tuberculosis* (*Mtb*) (27). Among the known β-lactamase inhibitors, avibactam strongly inhibits Bla_Mab_ (28) and reduces the MICs of various β-lactams against *Mab* (25, 29–31). Although clavulanate, tazobactam, and sulbactam are not potent inhibitors of Bla_Mab_, (27) clavulanate is the only orally-bioavailable agent and whether it exhibits synergy in combination with β-lactams against *Mab* is not sufficiently documented. Therefore, we have included avibactam and clavulanate in our study. Rifamycins were also included based on prior demonstration of synergy between carbapenems and rifamycins against *Mab in vitro* (32–34).

## RESULTS

### MICs of β-lactams against *M. abscessus*

We evaluated the antimicrobial activity of several β-lactam antibiotics, including nine cephalosporins (cefadroxil, cefprozil, cefuroxime, cefixime, ceftibuten, cefdinir, cefditoren, cefpodoxime, and cefoxitin), six carbapenems (ertapenem, meropenem, imipenem, doripenem, biapenem, and tebipenem), and a penem (faropenem) by determining their Minimum Inhibitory Concentrations (MIC) against *Mab* (Table 1). We preferentially tested oral cephalosporins that did not require more than twice daily dosing as their use in the clinical setting would be more convenient. MICs were also determined for three rifamycin antibiotics (rifabutin, rifapentine, and rifampin) and two β-lactamase inhibitors (clavulanate and avibactam), which were tested for synergy with β-lactams in subsequent experiments. The majority of cephalosporins exhibited high baseline MICs of 256 μg/mL, with the exception of cefoxitin and cefdinir at 64 μg/mL. The MICs of the carbapenems/penem were more variable, ranging from 8 μg/mL for imipenem to 256 μg/mL for ertapenem, tebipenem, and faropenem.

**Table 1.** Minimum inhibitory concentrations (MIC) of individual antibiotics tested against *M. abscessus* ATCC 19977 *in vitro*.

| β-lactam class | Drug | MIC ($\mu\text{g/mL}$) |
| --- | --- | --- |
| Cephalosporins | Cefadroxil | 256 |
|  | Cefprozil | 256 |
|  | Cefuroxime | 256 |
|  | Cefixime | 256 |
|  | Cefibuten | 256 |
|  | Cefdinir | 64 |
|  | Cefditoren | 256 |
|  | Cefpodoxime | 256 |
|  | Cefoxitin | 64 |
| Carbapenems | Ertapenem | 256 |
|  | Meropenem | 32 |
|  | Imipenem | 8 |
|  | Doripenem | 16 |
|  | Biapenem | 16 |
|  | Tebipenem | 256 |
| Penem | Faropenem | 256 |
| Rifamycins | Rifabutin | 32 |
|  | Rifapentine | 128 |
|  | Rifampin | 128 |
| β-lactamase inhibitors | Avibactam | 256 |
|  | Clavulanate | 256 |

### Several β-lactam combinations exhibit synergy against *M. abscessus*

A total of 206 paired combinations of antibiotics were initially screened for synergy via a condensed version of the checkerboard assay, as described in materials and methods. Combinations included all possible pairs of cephalosporins and carbapenems/penem, a rifamycin with either a cephalosporin or carbapenem/penem, and avibactam or clavulanate with a cephalosporin or carbapenem/penem (Figure 2).

**Figure 2.**
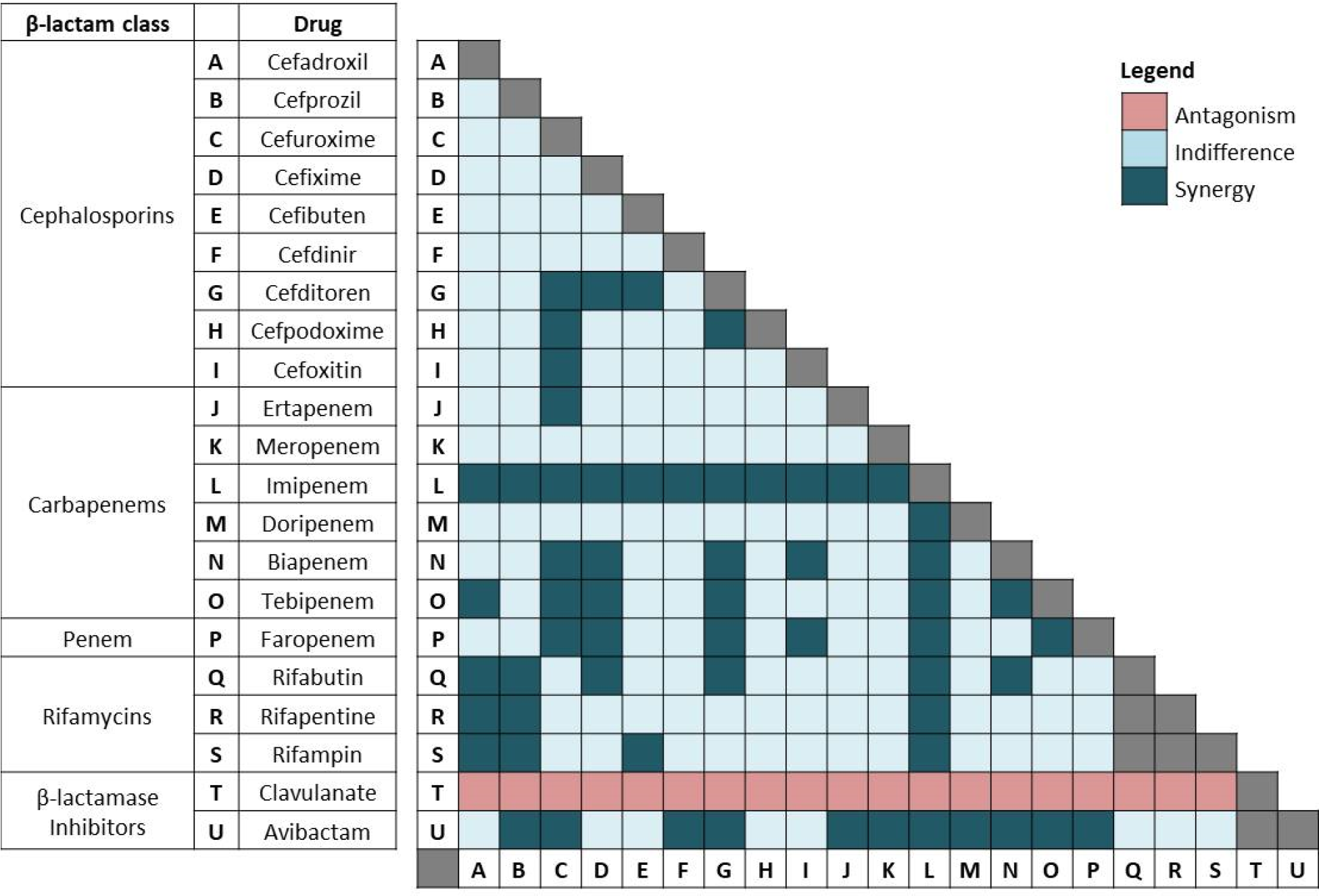
Results representing potencies of dual drug combinations against *M. abscessus* using checkerboard assay. Inhibition of *M. abscessus* growth in samples containing drug pairs at ¼ MIC or less of each drug is designated ‘Synergy’ and growth at ½ to 1x MIC of each drug is designated ‘Indifference.’ Growth of *M. abscessus* in samples containing drug pairs at 2× MIC of each drug is designated ‘Antagonism.’ These designations are based on the published guidelines for interpreting checkerboard assay results (35–37).

Of the initial 206 combinations screened, 60 combinations showed no growth at ¼ MIC or less for each drug and were further evaluated to verify synergy and determine the Fractional Inhibitory Concentration Index (FICI), using a full checkerboard titration assay as described in materials and methods (Figure 2). The FICI of a synergistic pair is a mathematical representation of the degree to which each drug contributes to synergy (35–37). Although substantial variability in the level of synergy among paired combinations was observed, there were a few notable trends. For example, imipenem was highly synergistic with all other drugs except clavulanate, and avibactam demonstrated synergy with all β-lactams except ceftibuten. Of note, all of the combinations that included clavulanate failed to inhibit growth of *Mab* in the presence of antibiotics at concentrations as high as 2× MIC. This antagonism was unexpected, but was reliably reproducible when the experiment was repeated.

A total of 24 out of the 60 combinations exhibited FICIs of ≤0.5 and thus were confirmed as synergistic (Table 2). To determine if the degree of synergy between paired combinations was sufficient to reduce MICs to within the therapeutic range, the Fractional Inhibitory Concentration (FIC) of each drug in combination was used to extrapolate expected MICs as a result of synergy. Although Clinical and Laboratory Standards Institute (CLSI) guidelines regarding MIC breakpoints against *Mab* are not currently available for most of the antibiotics tested, they have been established for cefoxitin and imipenem (38). Therefore, MIC breakpoints for all cephalosporins and carbapenems/penem were assumed to be the same as those for cefoxitin (≤16 μg/mL, susceptible; ≤64 μg/mL, intermediately susceptible) and imipenem (≤4 μg/mL, susceptible; ≤8 μg/mL, intermediately susceptible), respectively. With regard to the rifamycins, a breakpoint MIC of ≤1 μg/mL has been established for rifampin and rifabutin against *Mtb* and *Mycobacterium avium* complex (38) and the same principle was applied. Based on these presumed breakpoints, 5/24 combinations exhibited MICs within the fully susceptible range for both drugs, and an additional 7/24 combinations showed MICs that were considered moderately susceptible or intermediate. The remaining 12/24 combinations were unable to bring MICs below resistance breakpoints.

**Table 2.** Fractional inhibitory concentration indices (FICI) of synergistic β-lactam combinations. Combinations capable of reducing MICs to within susceptible/intermediate range based on established/presumed breakpoints are listed on the left. FICI and MICs in combination were extrapolated using data averaged from 2-3 replicate experiments.

| Drug combination | MIC of single drug (µg/mL) | MIC in combination (µg/mL) | FICI |
| --- | --- | --- | --- |
| Cefuroxime | 256 | 32 | 0.15 |
| Avibactam | 256 | 5 |  |
| Biapenem | 16 | 4 | 0.27 |
| Avibactam | 256 | 4 |  |
| Cefoxitin | 64 | 9 | 0.27 |
| Imipenem | 8 | 1 |  |
| Cefditoren | 256 | 26 | 0.27 |
| Imipenem | 8 | 1 |  |
| Imipenem | 8 | 2 | 0.30 |
| Doripenem | 16 | 2 |  |
| Cefuroxime | 256 | 33 | 0.30 |
| Cefditoren | 256 | 44 |  |
| Cefditoren | 256 | 26 | 0.32 |
| Biapenem | 16 | 4 |  |
| Cefdinir | 64 | 9 | 0.35 |
| Imipenem | 8 | 2 |  |
| Cefuroxime | 256 | 30 | 0.35 |
| Imipenem | 8 | 2 |  |
| Cefpodoxime | 256 | 28 | 0.36 |
| Imipenem | 8 | 2 |  |
| Imipenem | 8 | 2 | 0.42 |
| Biapenem | 16 | 3 |  |
| Imipenem | 8 | 2 | 0.47 |
| Avibactam | 256 | 64 |  |
| Tebipenem | 256 | 28 | 0.13 |
| Avibactam | 256 | 5 |  |
| Ertapenem | 256 | 64 | 0.27 |
| Avibactam | 256 | 4 |  |
| Ertapenem | 256 | 19 | 0.29 |
| Imipenem | 8 | 2 |  |
| Imipenem | 8 | 1 | 0.29 |
| Faropenem | 256 | 29 |  |
| Imipenem | 8 | 2 | 0.31 |
| Rifabutin | 32 | 3 |  |
| Cefadroxil | 256 | 40 | 0.34 |
| Tebipenem | 256 | 48 |  |
| Cefditoren | 256 | 64 | 0.38 |
| Tebipenem | 256 | 32 |  |
| Cefadroxil | 256 | 48 | 0.38 |
| Rifabutin | 32 | 6 |  |
| Cefadroxil | 256 | 64 | 0.38 |
| Rifapentine | 128 | 16 |  |
| Cefadroxil | 256 | 64 | 0.50 |
| Rifampin | 128 | 32 |  |
| Cefprozil | 256 | 64 | 0.50 |
| Rifabutin | 32 | 8 |  |
| Cefprozil | 256 | 64 | 0.50 |
| Rifapentine | 128 | 32 |  |

All of the 12 most synergistic combinations—those that that brought MICs within the therapeutic range—were pairs of either two β-lactams or a β-lactam and avibactam. Several of the remaining 12/24 combinations also exhibited a high degree of synergy based on FICI; however, their initial MICs were so high that the synergistic effect was insufficient to reduce MICs to below presumed breakpoints. We also observed a limit to the degree of synergy achievable with drugs exhibiting relatively low initial MICs, such as imipenem, biapenem, and doripenem. Combinations with these drugs resulted in up to a four-fold decrease in MIC, but this appeared to be the limit of reduction as MICs approached 2 μg/mL.

Additionally, the rifamycins showed synergy in combination with two of the earlier generation cephalosporins (cefprozil and cefadroxil); with at least a four-fold reduction in MIC for each combination. However, given the relatively low presumed MIC breakpoint of ≤ 1 μg/mL for rifamycins, none of the combinations brought MICs within the therapeutic range.

### β-lactam combinations reduce frequency of selection of spontaneous drug-resistant mutants

The 24 synergistic combinations were also evaluated to determine the frequency at which spontaneous resistant mutants are selected in the presence of paired drug combinations compared to each drug alone (Table 3). The frequency of resistant mutant selection was lower for all 24 combinations compared to the frequency for each drug individually. The greatest decrease was noted among the cephalosporins at >4 log reduction for six out of seven agents; with cefdinir being the exception, as the mutation frequency of this drug alone was lower than that of the other cephalosporins. Also of note, rifapentine and rifabutin exhibited the lowest frequency of resistant mutant selection, with no mutant colonies observed on any of the individual drug plates.

**Table 3.** Frequency of emergence of spontaneous drug-resistant mutants of *M. abscessus* when exposed to individual drugs and paired combinations that exhibit synergy *in vitro*.

| Drug combination | Resistance frequency |  | Drug combination | Resistance frequency |  |
| --- | --- | --- | --- | --- | --- |
|  | Individual drugs | Combination |  | Individual drugs | Combination |
| Cefuroxime<br>Avibactam | $> 2 \times 10^{-6}$<br>-- | $< 1 \times 10^{-10}$ | Tebipenem<br>Avibactam | $1.1 \times 10^{-7}$<br>-- | $< 1 \times 10^{-10}$ |
| Biapenem<br>Avibactam | $8.9 \times 10^{-8}$<br>-- | $3.3 \times 10^{-9}$ | Ertapenem<br>Avibactam | $8.3 \times 10^{-8}$<br>-- | $< 1 \times 10^{-10}$ |
| Cefoxitin<br>Imipenem | $> 2 \times 10^{-6}$<br>$1.9 \times 10^{-7}$ | $< 1 \times 10^{-10}$ | Ertapenem<br>Imipenem | $8.3 \times 10^{-8}$<br>$1.9 \times 10^{-7}$ | $4.3 \times 10^{-9}$ |
| Cefditoren<br>Imipenem | $> 2 \times 10^{-6}$<br>$1.9 \times 10^{-7}$ | $< 1 \times 10^{-10}$ | Imipenem<br>Faropenem | $1.9 \times 10^{-7}$<br>$< 1 \times 10^{-10}$ | $< 1 \times 10^{-10}$ |
| Imipenem<br>Doripenem | $1.9 \times 10^{-7}$<br>$9.9 \times 10^{-8}$ | $9.1 \times 10^{-9}$ | Imipenem<br>Rifabutin | $1.9 \times 10^{-7}$<br>$< 1 \times 10^{-10}$ | $< 1 \times 10^{-10}$ |
| Cefuroxime<br>Cefditoren | $> 2 \times 10^{-6}$<br>$> 2 \times 10^{-6}$ | $< 1 \times 10^{-10}$ | Cefadroxil<br>Tebipenem | $> 2 \times 10^{-6}$<br>$1.1 \times 10^{-7}$ | $1.3 \times 10^{-8}$ |
| Cefditoren<br>Biapenem | $> 2 \times 10^{-6}$<br>$8.9 \times 10^{-8}$ | $< 1 \times 10^{-10}$ | Cefditoren<br>Tebipenem | $> 2 \times 10^{-6}$<br>$1.1 \times 10^{-7}$ | $< 1 \times 10^{-10}$ |
| Cefdinir<br>Imipenem | $7.8 \times 10^{-9}$<br>$1.9 \times 10^{-7}$ | $< 1 \times 10^{-10}$ | Cefadroxil<br>Rifabutin | $> 2 \times 10^{-6}$<br>$< 1 \times 10^{-10}$ | $< 1 \times 10^{-10}$ |
| Cefuroxime<br>Imipenem | $> 2 \times 10^{-6}$<br>$1.9 \times 10^{-7}$ | $< 1 \times 10^{-10}$ | Cefadroxil<br>Rifapentine | $> 2 \times 10^{-6}$<br>$< 1 \times 10^{-10}$ | $< 1 \times 10^{-10}$ |
| Cefpodoxime<br>Imipenem | $> 2 \times 10^{-6}$<br>$1.9 \times 10^{-7}$ | $< 1 \times 10^{-10}$ | Cefadroxil<br>Rifampin | $> 2 \times 10^{-6}$<br>$> 2 \times 10^{-6}$ | $< 1 \times 10^{-10}$ |
| Imipenem<br>Biapenem | $1.9 \times 10^{-7}$<br>$8.9 \times 10^{-8}$ | $1.1 \times 10^{-9}$ | Cefprozil<br>Rifabutin | $> 2 \times 10^{-6}$<br>$< 1 \times 10^{-10}$ | $< 1 \times 10^{-10}$ |
| Imipenem<br>Avibactam | $1.9 \times 10^{-7}$<br>-- | $< 1 \times 10^{-10}$ | Cefprozil<br>Rifapentine | $> 2 \times 10^{-6}$<br>$< 1 \times 10^{-10}$ | $< 1 \times 10^{-10}$ |

## DISCUSSION

Given the growing prevalence of extensively drug-resistant *Mab* infections, development of novel treatment strategies is imperative. Although our current understanding of β-lactam targets in *Mab* is not comprehensive, we have leveraged the available data to begin developing synergistic treatment regimens with potential to treat *Mab* infections that are resistant to standard therapies.

In this study, we tested a total of 206 paired combinations of antibiotics *in vitro* against *Mab* reference strain ATCC 19977, 24 of which displayed synergy with FICI ≤ 0.5. Of these, 12 combinations achieved MIC reductions below presumed breakpoints for both drugs. Although a few studies have been published describing synergy between β-lactams and other antibiotic classes against *Mab in vitro* (32, 34, 39, 40), only one has assessed synergy between dual β-lactams (21). Our current study encompasses a broader array of combinations and offers a more comprehensive analysis of the synergistic activity of the two major β-lactam subclasses currently used to treat *Mab* infections; cephalosporins and carbapenems.

We hypothesize that the basis for synergy exhibited by β-lactam pairs is their non-redundant selective inhibition of distinct transpeptidases that synthesize PG in *Mab*. If only one type of enzyme existed as the target for β-lactams, the pairs would likely exhibit additive activity rather than synergy. Variation in the level of inhibition of L,D-transpeptidases of *Mab* (21) and *Mtb* (20, 24, 41) by agents of this antibiotic class supports this hypothesis. We further hypothesize that, at the molecular level, the structures of β-lactams that exhibit synergy against *Mab* most effectively complement the structure of the binding sites available in the transpeptidases of this organism; thereby favoring initial binding and subsequent interaction to bring about effective inhibition of the enzymes. Differences in binding affinity and kinetics of inhibition by different β-lactams against LDTs of *Enterococcus faecium* (42) illustrate the basis for this hypothesis.

Five out of 24 synergistic combinations included avibactam. In these combinations, the MICs of three β-lactams (cefuroxime, imipenem, and biapenem) were reduced to below therapeutic breakpoints. Therefore, avibactam appears to be a viable adjunct to β-lactam-based treatment regimens. However, its current coformulation with ceftazidime (which itself does not exhibit valuable activity against *Mab* (28, 30)) and intravenous administration limit its usefulness. Relevant to the strategy of combining a carbapenem with a β-lactamase inhibitor against *Mab*, several coformulated agents have recently been developed. These include FDA-approved meropenem-vaborbactam (Melinta Therapeutics) and imipenem-relebactam (Merck), which recently completed phase III trials. Coformulations of cefepime-zidebactam (Wockhardt) and meropenem-nacubactam (Roche) are currently in phase II of development. Although these compounds have yet to be tested against *Mab*, our data suggest they could be viable options for treatment of *Mab* infections and would potentially simplify β-lactam-based regimens. Interestingly, clavulanate was found to be antagonistic in combination with all β-lactams tested in this study, although the mechanism for this is unclear. As there is no precedent for antagonism of β-lactam and clavulanate combinations against other bacteria, we speculate the following hypotheses. Clavulanate is metabolized by *Mab* and the resulting metabolite alters the rate of influx or efflux of β-lactams, thereby reducing the effective concentration available to bind target enzymes. It is also possible that binding of the clavulanate metabolite to target enzymes may alter their binding kinetics to β-lactams. In addition, the possibility of the clavulanate metabolite directly competing for binding sites associated with the 4-carbon core ring of β-lactams cannot be ruled out. Further study would be necessary to elucidate the underlying mechanism for this phenomenon, including possibilities not considered above.

Another measure of synergy is the frequency of selection of spontaneous resistant mutants. For a synergistic pair, the frequency of resistant mutant selection would ideally be lower than the product of frequencies associated with either drug alone. The majority of drug pairs with the lowest FICIs selected resistant mutants with a frequency of <1 × 10^−10^, which approaches the product of the individual drugs (Table 3). However, due to physical limitations of the number of *Mab* CFU that could be used to identify resistant mutants, we were unable to obtain the exact frequency of mutant selection for several paired combinations. Based on these observations, we propose that the majority of pairs identified here exhibit synergy in both antimicrobial activity and reduction of selection of drug-resistant mutants.

Although significant variability in MIC exists among *Mab* strains and *in vitro* drug susceptibility data does not always correlate with clinical efficacy (43), the novel β-lactam combinations identified here using the reference *Mab* strain ATCC 19977 could be leveraged for further preclinical assessment, including *in vivo* efficacy against drug-resistant clinical isolates. As *Mab* treatment generally necessitates a regimen consisting of at least 3-4 agents, the addition of other antibiotic classes or β-lactamase inhibitors to these synergistic β-lactam combinations may further potentiate MIC reduction and improve efficacy. This may also allow clinicians to avoid use of the more cytotoxic antibiotics such as amikacin, especially in the context of prior adverse effects.

Based on current trends of *Mab* strains isolated in clinics, resistance to an increasing number of drugs is likely to continue over the next several years; further compromising our ability to treat disease resulting from this pathogen. New antibiotics and coformulations are currently in development, but none are primarily intended for treatment of *Mab* or other NTM infections and it could take several years for efficacy studies to be completed. Repurposing of currently-available antibiotics in novel combinations, such as those identified here, may provide vital immediate therapeutic options for patients failing standard *Mab* treatment regimens and facilitate rapid implementation in the clinical setting.

## MATERIALS AND METHODS

### Bacterial strains and *in vitro* growth conditions

The *Mab* reference strain ATCC 19977 (ATCC, Manassas, VA) was used for all experiments. Strains were grown in Middlebrook 7H9 broth (Difco) supplemented with 0.5% glycerol, 10% albumin-dextrose-catalase enrichment, and 0.05% Tween80, at 37°C with constant shaking at 220 RPM in an orbital shaker. All drugs were obtained from the following commercial vendors: ertapenem (Toronto Research Chemicals), rifampin, meropenem, imipenem, doripenem, biapenem, faropenem, tebipenem, and all cephalosporins (Sigma-Aldrich). To assess the quality of these compounds, a few were randomly selected and assessed by liquid chromatography-mass spectrometry. The purity of compounds ranged from 95% to 99%.

### Minimum Inhibitory Concentration

Minimum inhibitory concentration (MIC) of each drug against *Mab* was determined using the standard broth dilution method (44, 45) in accordance with CLSI guidelines specific for this organism (38). In summary, powdered drug stocks were reconstituted in dimethyl sulfoxide (DMSO) and two-fold serial dilutions were prepared in Middlebrook 7H9 broth to obtain final drug concentrations ranging from 256 μg/mL to 1 μg/mL in 96-well plates in a final volume of 200 μL. 10^5^ colony forming units (CFU) of *Mab* from exponentially growing culture was added to each well. *Mab* culture without drug and 7H9 broth alone were included in each plate as positive and negative controls, respectively. Plates were incubated at 30°C for 72 hours per CLSI guidelines. MIC was assessed via visual inspection to determine growth or lack thereof and an MIC for each drug was recorded as the lowest concentration at which *Mab* growth was not observed. All MIC assessments were repeated to verify results.

### Checkerboard Titration Assay

The checkerboard titration assay is a modified broth dilution assay and was performed as previously described (35–37). An initial synergy screen of two-drug combinations was performed at four different concentrations based on each drugs’ respective MIC: 2× MIC, MIC, ½ MIC, and ¼ MIC for each drug in combination via two-fold serial dilutions in a 96-well plate. 10^5^ CFU of *Mab* from exponentially growing culture were inoculated into each well, with positive and negative controls as described above for the MIC assay, and plates were incubated at 30°C for 72 hours. Plates were visually inspected for *Mab* growth or lack thereof. Drug combinations that inhibited *Mab* growth at ¼ MIC or less of each drug were considered to have some degree of synergy and were chosen for additional synergy testing.

To confirm the degree of synergy, two drugs were added to Middlebrook 7H9 broth in a 96-well plate, each starting at 2× MIC and serially diluted two-fold up to 1/32× MIC, so all possible two-fold dilution combinations from 2× to 1/32× MIC were assayed. 10^5^ CFU of *Mab* were inoculated into each well. Plates were incubated at 30°C and evaluated for *Mab* growth by visual inspection at 72 hours. The Fractional Inhibitory Concentration (FIC) of each drug in combination was determined as described previously (35–37). The FIC of a drug in a sample is calculated as the concentration of the drug divided by the MIC of the drug when used alone. The FIC Index (FICI) is the sum of the FIC of two drugs in a sample. The FICI was calculated for each combination of drugs that inhibited *Mab* growth at less than ½ the MIC of each individual drug. An FICI of ≤0.5 was interpreted as synergy, >0.5 to 2 as indifference, and >2 as antagonism. As an internal control, the MIC of each individual drug was also assessed via broth microdilution within each plate. All combinations with an FICI of ≤0.5 were tested in triplicate to confirm reproducibility and an average FICI was calculated and reported here.

### Determination of frequency of spontaneous drug resistance emergence

Any drug combination with an FICI of ≤0.5 was further assessed for frequency of spontaneous drug resistance. The CFU/mL of *Mab* in culture at an A_600nm_ of 1.0 was initially determined as follows. *Mab* was grown to exponential phase, adjusted to an A_600nm_ of 1.0 in Middlebrook 7H9 broth and was serially diluted 10-fold in this broth. 100 μL of each dilution was plated onto Middlebrook 7H10 agar, which were incubated at 37°C for 72 hours. Resultant CFU counts were used to determine *Mab* CFU density in culture. This assessment was repeated three times and the mean *Mab* CFU density was used in calculations in subsequent experiments.

To determine frequency of spontaneous drug resistance emergence, 10 mL of *Mab* culture grown to exponential phase in 7H9 broth was used to prepare a suspension at A_600nm_ of 1.0, and 1.0 mL of this suspension was inoculated onto each of 10 total Middlebrook 7H10 agar plates, which were supplemented with either a single drug or a combination of two drugs. These assessments were performed at the MIC for single-drug and combination plates, to promote selection of resistant mutants. CFU were counted after 7 days of incubation at 37°C. The frequency of drug-resistant mutants was determined from the number of spontaneous mutants observed as a percentage of the input CFU inoculum.

## ACKNOWLEDGMENTS

This work was supported by the Cystic Fibrosis Foundation award LAMICH17GO and NIH award R21 AI121805. ESR was supported by NIH T32 AI007291. The content is solely the responsibility of the authors and does not necessarily represent the official views of the Cystic Fibrosis Foundation or the National Institutes of Health.

